# Tooth development in frogs: Implications for the re-evolution of lost mandibular teeth and the origin of a morphological innovation

**DOI:** 10.1101/2025.02.17.638649

**Authors:** Daniel J. Paluh, Madeline Brinkman, Kyliah Gilliam-Beale, Daniela Salcedo-Recio, Jacob Szafranski, James Hanken, Gareth J. Fraser

**Affiliations:** Department of Biology, University of Dayton, Dayton, OH USA 45469; Museum of Comparative Zoology, Harvard University, Cambridge, MA USA 02138; Department of Biology, University of Florida, Gainesville, FL USA 32611

**Keywords:** Anura, Tadpole, Odontogenesis, Dollo’s Law, Novelty

## Abstract

Teeth have been a prominent feature of most vertebrates for 400 million years, and the core regulatory network underlying embryonic tooth formation is deeply conserved from fishes to amniotes. In frogs, however, odontogenesis is delayed, occurring instead during the postembryonic metamorphosis. Nearly all adult frogs have teeth on the upper jaw and lack lower jaw dentition, but a single species re-evolved mandibular teeth. Developmental-genetic mechanisms that underlie tooth formation in frogs are poorly understood, including whether an ancestral program is partially retained in the lower jaw that could facilitate the evolutionary reappearance of lost mandibular teeth. Using a developmental series of an unconventional model species, we assessed 1) if the gene network underlying odontogenic competence is conserved in the late-forming teeth of frogs; 2) if unique keratinized mouthparts, which function as an alternative feeding tool in anuran larvae, impede tooth induction; and 3) if transient tooth rudiments form in the anuran mandible. The tooth development network is conserved in the frog upper jaw, which displays dental expression patterns comparable to those of other vertebrates. There is, however, no evidence of tooth development in the mandible. Teeth emerge before keratinized mouthparts degenerate, but their location may be spatially constrained by keratin, and gene expression patterns of keratinized mouthparts and teeth overlap. We hypothesize that the novel mouthparts of tadpoles did not arise *de novo* but instead originated by partially co-opting the developmental program that typically mediates true tooth development.

## INTRODUCTION

Teeth are complex, mineralized structures composed of a pulp cavity, dentin, and enamel. They originated in stem gnathostomes over 400 million years ago. Because teeth play a crucial role in the acquisition and processing of food, they are broadly conserved among living chondrichthyans, actinopterygians, and sarcopterygians, although their shape, size, location, and number vary widely due to broad variation in feeding mechanisms. They are rarely lost. The developmental genetics of tooth formation has been studied extensively in chondrichthyans, teleosts, and amniotes, and the highly complex core gene regulatory network (GRN) underlying odontogenesis is deeply conserved [1–3]. Tooth development in embryonic jaws of fishes, salamanders, and most amniotes begins with formation of an odontogenic band (OB), which expresses the genes *Shh*, *Pitx2*, and *Sox2* [1, 3–6] and defines the region of oral epithelium that is competent to form teeth. Yet, compared to the rich knowledge of odontogenesis in most vertebrate clades, the developmental genetics of tooth formation in frogs is very poorly understood.

Among living amphibians, all salamanders and caecilians possess teeth on both upper and lower jaws, but nearly all frogs lack lower jaw (i.e., mandibular) teeth and variably possess teeth on the upper jaw and palate [7]. The only published study on the developmental genetics of tooth induction in anurans failed to document an OB marked by *Shh* expression before tooth bud formation in the upper jaw of *Xenopus tropicalis* [8]. This suggests that the regulatory network underlying tooth development may be considerably modified in frogs, perhaps due to the delayed onset of odontogenesis. Frog teeth form during metamorphosis [9], unlike other vertebrates in which teeth develop in the embryo.

Delayed onset of tooth development in frogs is a likely consequence of the unique keratinized mouthparts of tadpoles because a non-keratinized oral epithelium is thought to be required for initiation of odontogenesis [10–11]. In most tadpoles, the mouth is surrounded by keratinized jaw sheaths (i.e., horny beak), several rows of keratodonts (i.e., keratinized labial “teeth”), and sensory papillae on the upper and lower labium of the oral disk. The jaw sheaths and keratodonts are composed of several epidermal cell types stacked into a column [12–14]. The basal cells are proliferative, enabling continuous replacement of these structures [15–16]. The tadpole feeding apparatus is considered an evolutionary novelty and key innovation [17] that enables frog larvae to feed on plant material and detritus. The evolutionary origin of the tadpole feeding apparatus, however, is entirely unknown [18], as are the developmental genetics that mediate its formation. *Xenopus*, a popular anuran model organism, lacks these keratinized structures, which has impeded their study.

Of the nearly 8,000 species of living frogs [19], only Guenther’s marsupial frog, *Gastrotheca guentheri*, possesses teeth on the dentary bone of the lower jaw. Mandibular teeth were lost in the common ancestor of frogs more than 200 million years ago [20], but they were later regained in *G. guentheri* or one of its direct ancestors during the Miocene epoch [21–22]. The presence of mandibular teeth in this species is a notable exception to Dollo’s law of irreversibility, which holds that complex structures lost over evolutionary time cannot be regained in the same form [23]. Yet, evidence of this instance of trait reversal is derived solely from phylogenetic reconstruction of character evolution; its developmental basis is entirely unknown. Evolutionarily lost traits are rarely entirely absent from the developmental program of descendants [24], and the presence of transient, vestigial organs can preserve genetic and developmental pathways that may provide a mechanism for the reacquisition of lost phenotypes [25–28]. It is thus possible that an ancient tooth development program is initially expressed in the lower jaw of many living frogs, forming a transient OB and tooth bud rudiments before being disrupted by a conserved mechanism that originated in proto-frogs.

In this study, we characterize development of first-generation teeth in the Cuban tree frog, *Osteopilus septentrionalis*, which has a typical tadpole with keratinized mouthparts and an adult with teeth on the upper jaw but a toothless mandible (Fig. 1). We evaluate histological anatomy and expression of several key genes across a comprehensive developmental series to 1) determine the precise timing of odontogenesis in the upper jaw, 2) assess if an OB forms comparable to those in other toothed vertebrates, 3) look for evidence of inductive or inhibitory interactions between the keratinized mouthparts of tadpoles and developing teeth, and 4) evaluate if tooth development is initiated on the lower jaw but then abandoned.

**Fig. 1.**
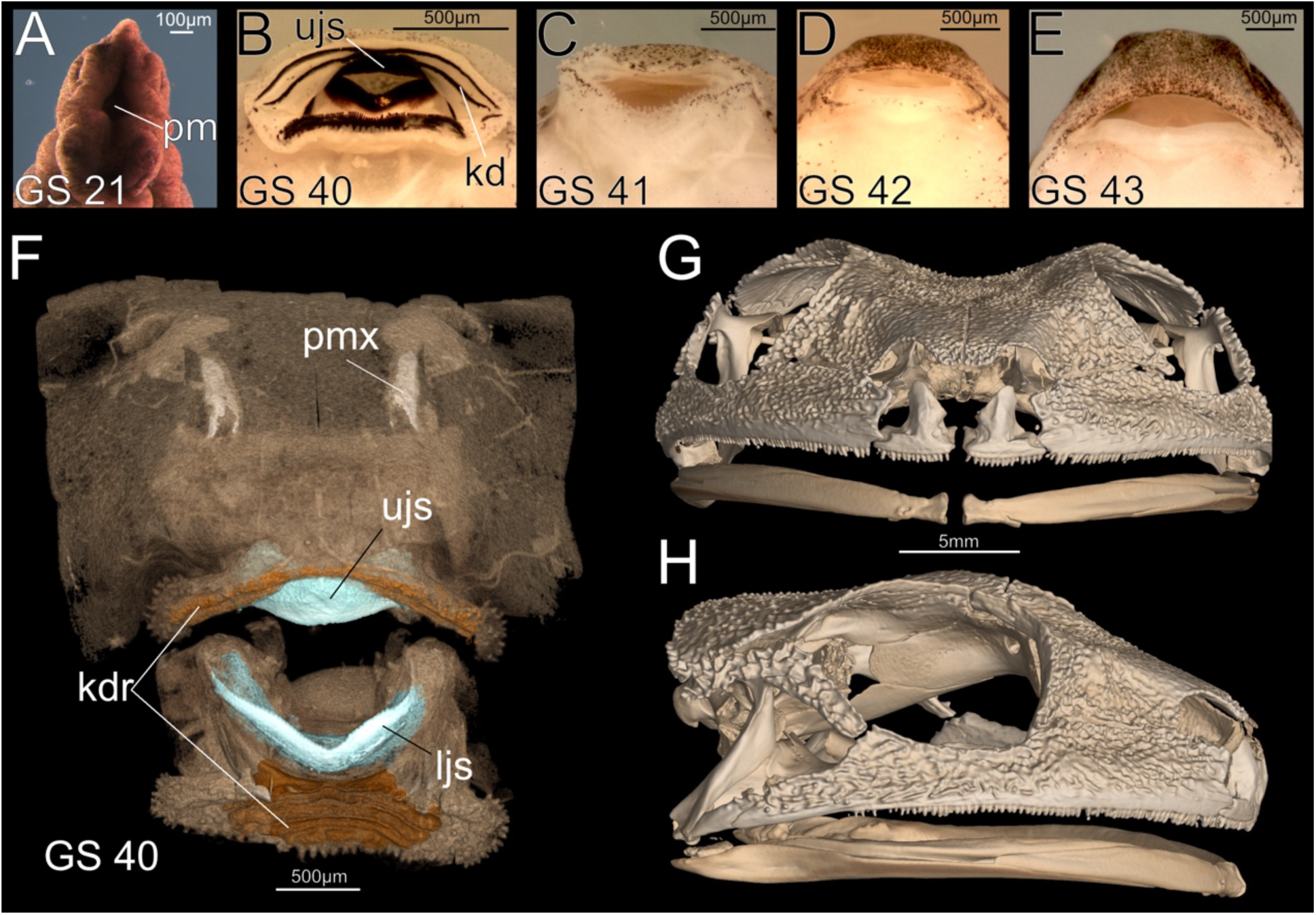
Anatomy and development of the feeding apparatus in the Cuban tree frog, *Osteopilus septentrionalis*. A–E, Progressive development of the mouth: stomodeum (primary embryonic mouth; A), tadpole oral disk with keratinized mouthparts (B), atrophy of keratinized mouthparts (C), and formation of post-metamorphic frog jaws (D, E). F–H, microCT scans of tadpole feeding apparatus (F) and adult skull with a toothed upper jaw and edendate mandible (G and H). Abbreviations: kd, keratodonts; kdr, keratodont ridges; ljs, lower jaw sheath; pm, primary mouth; pmx, premaxilla; ujs, upper jaw sheath. See SI Methods for scanning methods.

## RESULTS

### Establishing odontogenic competence on the upper jaw

In *Osteopilus septentrionalis*, Gosner stage (GS) 40 can be considered the final “tadpole” stage; the larval feeding apparatus, keratinized mouthparts, and vent tube are still present, but hindfoot tubercles have begun to form (Fig. 1B and F). We evaluated oral histology in seven GS 40 tadpoles, four of which show no evidence of tooth development. The oral mucosa of the buccal roof is composed of stratified squamous epithelium with a cornified outermost layer, the stratum corneum, that is contiguous with the keratinized upper jaw sheath of the tadpole feeding apparatus (Fig. 2A, black cornified layers). Loose, uncondensed mesenchyme is present between the oral epithelium and the suprarostral cartilage of the upper jaw. The remaining three specimens show the earliest histological indication of tooth initiation on the upper jaw. Cells of the basal layer of the oral epithelium are transitioning from a squamous to a columnar shape, which produces a localized epithelial thickening located at the posterior extent of the keratinized jaw sheath cells (Fig. 2B). Underlying mesenchyme is beginning to condense, and abundant dividing cells (i.e., mitotic figures) within the oral epithelium suggest high cell proliferation rates.

**Fig. 2.**
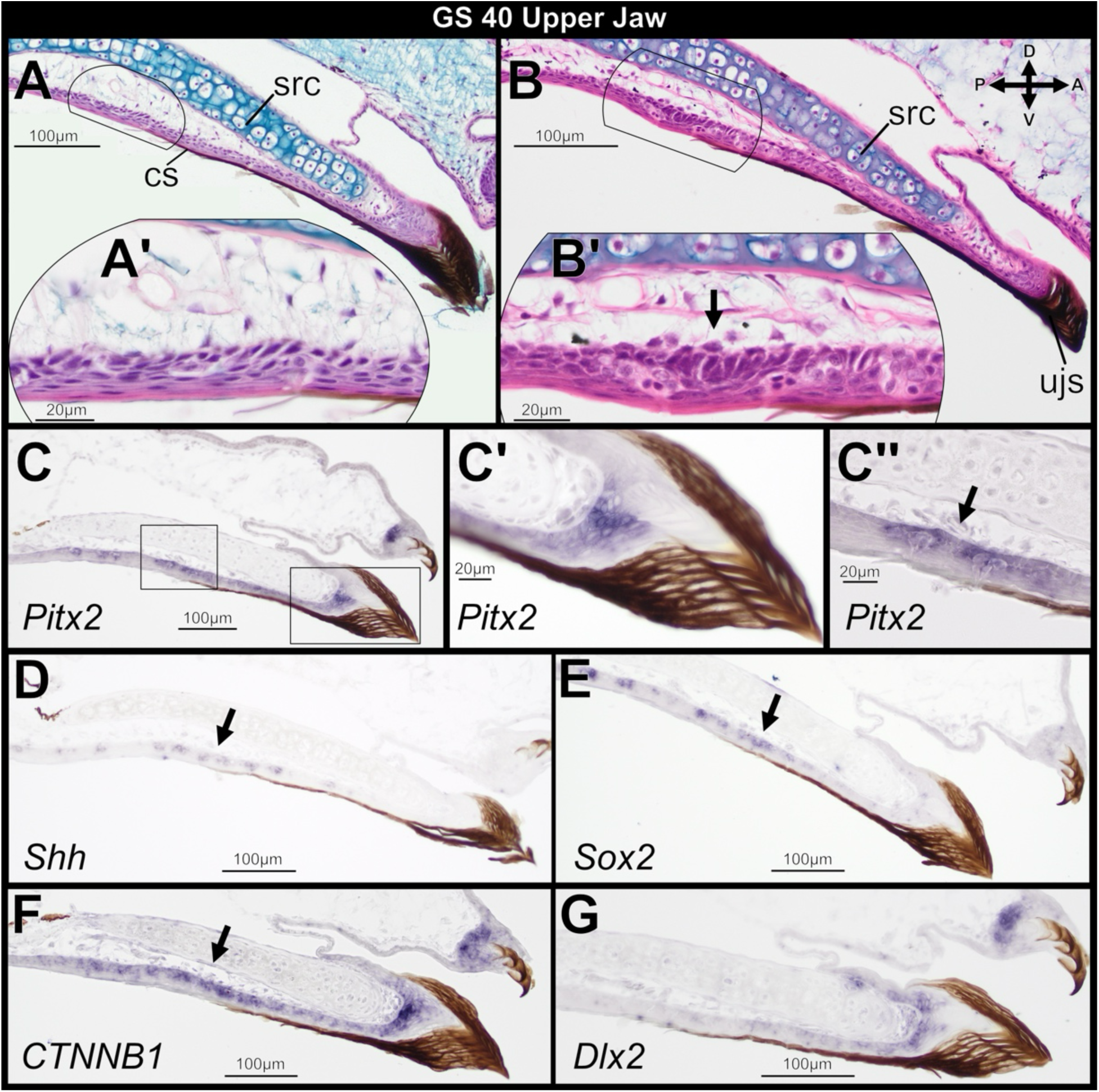
Sagittal sections through the larval upper jaw of *Osteopilus septentrionalis,* Gosner stage 40, stained with H&E (A, B) and *in situ* hybridization (C–G). The earliest histological indication of tooth induction occurs with the formation of a localized epithelial thickening (arrow in B’). *Pitx2*, *Shh*, *Sox2*, and *CTNNB1* mark the zone of upper jaw oral epithelium where tooth buds will form (arrows in C”, D, E, and F). *Pitx2*, *CTNNB1*, and *Dlx2* are also highly expressed in the tadpole jaw sheath and keratodonts (C’, F, and G). Insets (C’ and C”) show higher magnification detail of regions outlined in C. Abbreviations: cs, cornified sheath cells; src, suprarostral cartilage. Orientation axes: D, dorsal; V, ventral; A, anterior; P, posterior.

Expression of several genes known to mediate tooth initiation accompanies the above histological changes. *Pitx2* is the earliest epithelial marker of tooth competence known in tetrapods [29]. At GS 40, the gene is expressed before epithelial thickening in the upper jaw oral epithelium where tooth buds will form (Fig. 2C and 2C’’). Unexpectedly, *Pitx2* also is highly expressed within epithelial basal cells that support the upper jaw sheath and keratodonts of the keratinized tadpole feeding apparatus (Fig. 2C and 2C’). This expression is asymmetric—lingually in the jaw sheath and labially in the keratodonts. *Pitx2* expression is continuous between the epithelial base of the upper jaw sheath and the region of the upper jaw oral epithelium where tooth buds will form. *Shh* is weakly expressed in the upper jaw oral epithelium before tooth development but is absent in the epithelial cells that support the keratinized jaw sheath and keratodonts (Fig. 2D). *Sox2* is expressed in the oral epithelium preceding tooth development and is also weakly expressed in the epithelial cells at the base of the keratodonts (Fig. 2E). *CTNNB1*, which encodes for β-Catenin protein, is strongly expressed in the upper jaw oral epithelium, the upper jaw sheath, and keratodonts (Fig. 2F). Unlike *Pitx2*, *CTNNB1* is more widely expressed throughout the basal cells of the jaw sheath and keratodonts, as well as underlying mesenchyme. Finally, *Dlx2* is expressed within the epithelial basal cells and underlying mesenchyme of the keratinized jaw sheath and keratodonts (Fig. 2G). *Dlx2* expression within the oral epithelium is limited to the anterior region that underlies the jaw sheath; unlike *Pitx2* and *CTNNB1*, it does not extend posteriorly to where the localized epithelial thickenings will form.

### Emergence of the dental lamina and first-generation teeth on the upper jaw

Gosner stage 41 is characterized by atrophy of the keratinized mouthparts and vent tube (Fig. 1C). Of the nine specimens sectioned at this stage, six retain keratinized mouthparts while three are undergoing mouthpart atrophy. Odontogenesis has begun in the upper jaw of all specimens. Tooth buds first appear ventral to distal and midshaft regions of the suprarostral cartilages in parasagittal sections; there is no indication of tooth development in midline sections where these paired cartilages meet. The earliest stage of dental lamina emergence and tooth placode formation is visible: labial and lingual edges of the columnar basal cells that delimit the oral epithelial thickening are invaginating into the underlying ectomesenchyme, forming a concave plate of cells (Fig. 3A; described as an inverted trough in *Xenopus* [30]). The ectomesenchyme is condensing and protrudes apically into the center of the thickened epithelium. The dental lamina has become asymmetric, with the labial edge of the epithelial plate invaginating more deeply and in a posterior direction, forming a roof and differentiating into the dental epithelium that encloses condensed mesenchymal cells that will form the dental papilla (Fig. 3B and 3C). These first-generation tooth buds develop superficially, near the surface of the oral mucosa, similar to dental lamina emergence in *Xenopus* [31]. The bicuspid crown pattern has already emerged in tooth buds of several specimens that retain the jaw sheath (Fig. 3D). The jaws undergo restructuring as the tadpole mouthparts degenerate, but the developmental stage of first-generation tooth buds does not advance dramatically during this time. Finally, the genes *Pitx2*, *Shh*, Sox2, *CTNNB1*, and *Dlx2* are expressed in the emerging dental lamina of the upper jaw (Figs. 3E–J). Expression of *Pitx2*, *Sox2*, *CTNNB1*, and *Dlx2* is no longer detectable in epithelial cells that support the keratinized jaw sheath, which is beginning to degenerate. *Pitx2* is widely expressed within oral epithelium on the roof of the mouth, in effect connecting the emerging dental lamina that will give rise to tooth buds on the palate (vomerine teeth) and the dental lamina that will give rise to teeth on the upper jaw (Fig. 3F). *Dlx2* is expressed in epithelial cells delimiting the dental lamina, as well as in condensed mesenchyme that will differentiate into the dental papilla (Fig. 3J).

**Fig. 3.**
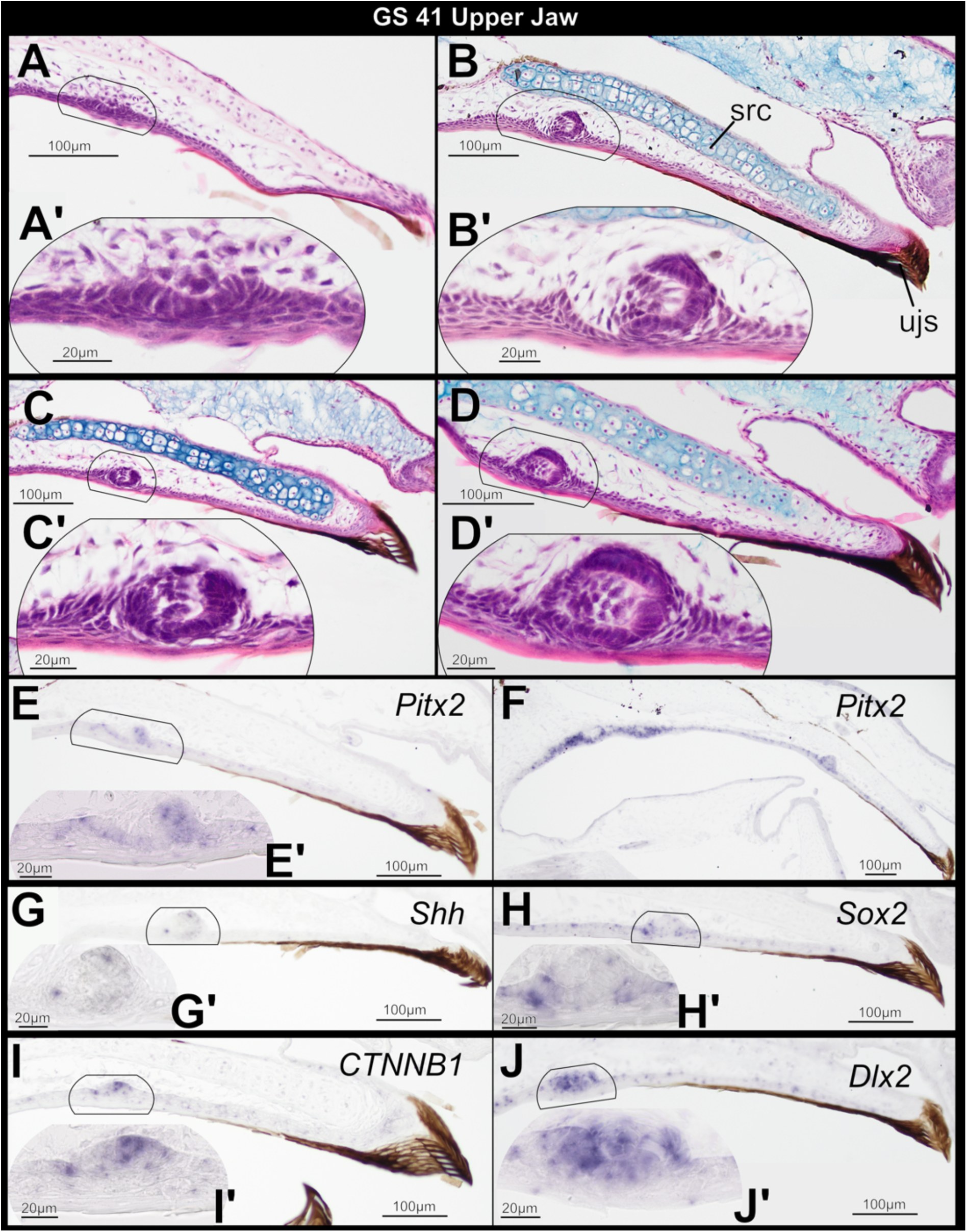
Sagittal sections through the larval upper jaw of *Osteopilus septentrionalis,* Gosner stage 41, stained with H&E (A–D) and *in situ* hybridization (E–J). The dental lamina emerges (A) and tooth placodes form (B–D) coincident with keratinized mouthparts. *Pitx2*, *Shh*, *Sox2*, *CTNNB1*, and *Dlx2* are expressed in emerging tooth placodes (E–J).

Gosner stage 42 is characterized by emergence of fully formed forelimbs. The simple and narrow mouth opens anterior to the external nares (Fig. 1D), and there are remnants of the tadpole oral disk structures. Teeth in several stages of development are visible along the upper jaw row; younger tooth buds are located nearer the midline. Tooth buds have reached the cap stage of morphogenesis, and the epithelial plate has invaginated more deeply into the ectomesenchyme (Fig. 4A–D). The dental epithelium has differentiated into inner and outer layers, and condensed ectomesenchymal cells have formed the dental papilla. A prominent successional dental lamina, which gives rise to future tooth generations, has emerged lingual to the first-generation teeth and connects the row of emerging tooth buds (Figs. 4B, S1). In a subset of tooth buds, cells of the inner dental epithelium have differentiated into ameloblasts and dental papilla cells have differentiated into odontoblasts. The first signs of tooth matrix secretion by odontoblasts are visible (Fig. 4C and D). At later metamorphic stages (GS 43–46), the jaws rapidly extend posteriorly (Fig. 1E), bringing the jaw joint posterior to the eye, while first-generation tooth buds continue to undergo dentinogenesis. The latter also migrate toward the ossifying maxillary and premaxillary bones, but they are not functional as they are yet to implant in bone or erupt through the oral mucosa (Fig. 4E–H). The successional dental lamina continues to invaginate more deeply into underlying ectomesenchyme and assumes a cup shape at its extremity (Fig. 4G). At the same time, condensing ectomesenchyme surrounds the extremity of the successional dental lamina (Fig. 4G and H), indicating that differentiation of second-generation tooth buds begins before metamorphosis is complete. First-generation teeth in frogs are non-pedicellate [32], and we saw no indication of pedicel formation.

**Fig. 4.**
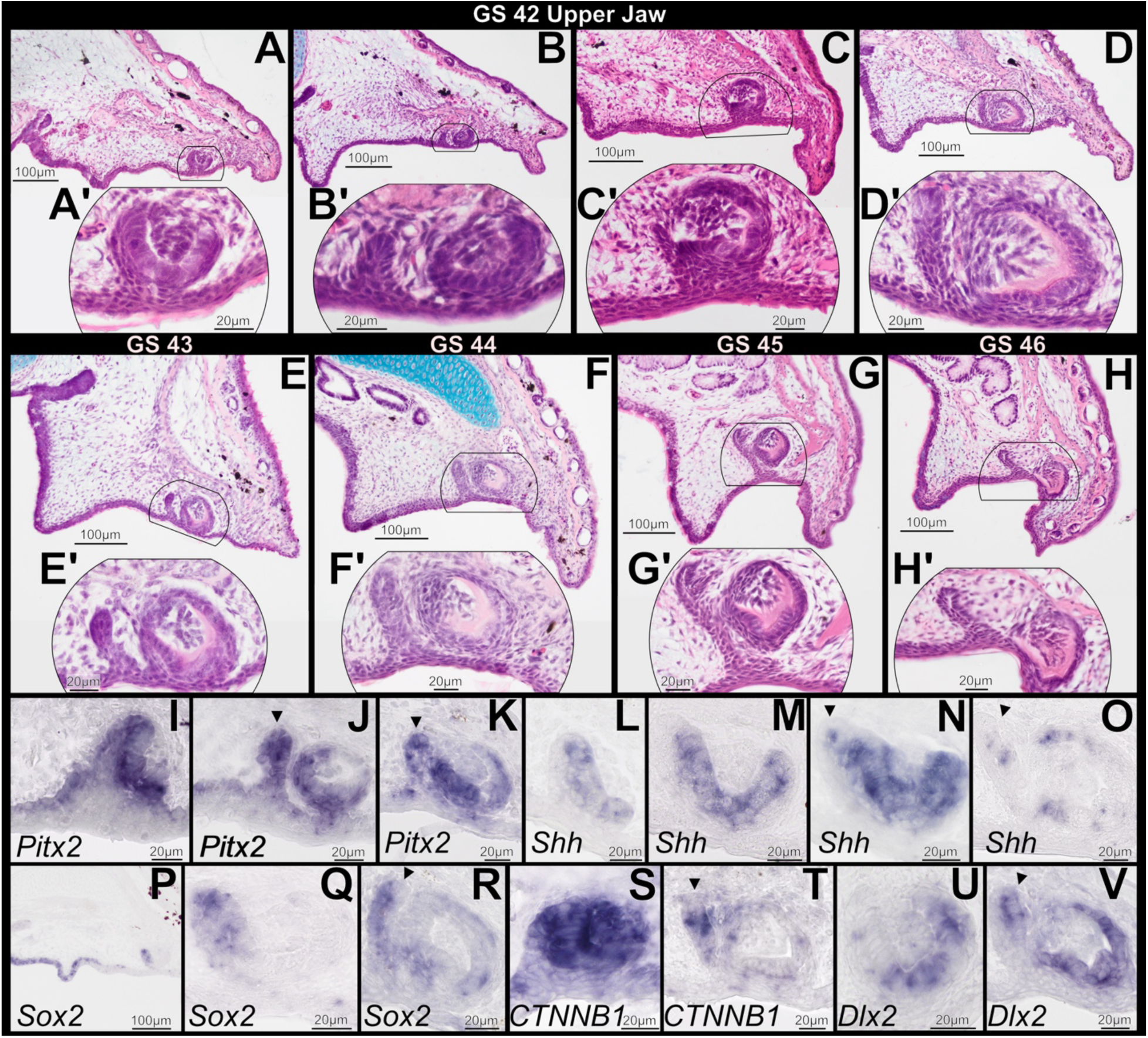
Sagittal sections through the upper jaw of *Osteopilus septentrionalis* during and immediately following metamorphosis, Gosner stages 42–46, stained with H&E (A–H) and *in situ* hybridization (I–V). Tooth buds at GS 42 have attained the cap stage of morphogenesis and a successional dental lamina has emerged (A–D). The first signs of tooth matrix secretion by odontoblasts are also visible (C, D). Developing tooth buds do not become functional until after the completion of metamorphosis (E–H). *Pitx2*, *Shh*, *Sox2*, *CTNNB1*, and *Dlx2* are expressed in dental tissues throughout morphogenesis (I–V). Arrowheads denote expression in the successional dental lamina.

During GS 42–46, the genes *Pitx2*, *Shh, Sox2*, *CTNNB1*, and *Dlx2* are expressed in developing first-generation teeth (Fig. 4I–V). *Pitx2* is expressed in the inner and outer dental epithelia of tooth buds, as well as the emerging successional dental lamina (Fig. 4I–K). *Shh* is expressed within the inner dental epithelium of tooth buds at the cap stage (Fig. 4M and N) and also marks the free, distal end of the successional dental lamina (Fig. 4N and O). At GS 42, *Sox2* is strongly expressed in olfactory and upper jaw oral epithelium posterior to the tooth buds, as well as in the lingual portion of the dental epithelium (Fig. 4P and Q). *Sox2* is also expressed in the emerging successional dental lamina (Fig. 4R). *CTNNB1* is strongly expressed in the dental epithelium and dental papilla during the cap stage (Fig. 4S). It also is present in the successional dental lamina and condensing ectomesenchyme at later stages (Fig. 4T). At GS 42, *Dlx2* expression is limited to the labial portion of the dental epithelium, but it later expands to the entire dental epithelium as well as the free, distolabial end of the successional dental lamina (Fig. 4U and V).

### Evaluating odontogenic potential on the anuran lower jaw

We examined the lower jaw in all 45 sectioned specimens for any histological evidence of odontogenesis in the mandible coincident with tooth formation in the upper jaw. The tadpole lower jaw has a complex shape and is segmented into two parts: anteromedially, a V-shaped infrarostral cartilage supports the keratinized lower jaw sheath, while posteriorly and laterally, a pair of transversely oriented Meckel’s cartilages each articulate with the infrarostral via an intramandibular commissure [17]. During metamorphosis, infrarostral and Meckel’s cartilages fuse to form a single rod of lower jaw cartilage.

Parasagittal sections through articulating Meckel’s and infrarostral cartilages at GS 40 reveal a simple, squamous epithelium overlying the lingual side of Meckel’s cartilage (Fig. 5A). In this same plane, a stratified, squamous epithelium that is contiguous with the lower jaw sheath overlies the labial side of the infrarostral cartilage (Fig. 5B). The histology of this epithelium resembles the oral epithelium of the upper jaw before odontogenesis (Fig. 2A). In sections closer to the midline, most of the oral epithelium overlying the infrarostral cartilage is simple and squamous (Fig. 5B). In contrast, epithelia associated with the keratinized lower jaw sheath is complex, including columnar basal keratinocytes, several layers of stratified sheath cells that form the lingual and labial surfaces of the jaw sheath, and a central column of proliferating cone cells, which form the cutting edge of the jaw (Fig. 5C). None of the GS 40 specimens shows any histological evidence of a localized epithelial thickening overlying Meckel’s or infrarostral cartilage, nor is there any other indication of odontogenesis in the lower jaw at this stage. However, gene expression patterns within the keratinized mouthparts of the lower jaw are comparable to those in the upper jaw at the same stage. *Pitx2*, *CTNNB1*, and *Dlx2* are highly expressed in the lower jaw sheath and keratodonts (Fig. 5C, E, and F). *Pitx2* expression is asymmetric—lingually in the jaw sheath but labially in the keratodonts—while *CTNNB1* and *Dlx2* expression is more widespread in the basal and suprabasal epithelial cells that support these structures. Expression of these markers is limited to epithelial cells associated with the keratinized lower jaw sheath and does not extend posteriorly to the oral epithelium that overlies the infrastrostral cartilage, a pattern that differs markedly from *Pitx2* and *CTNNB1* patterns in the upper jaw. *Sox2* is weakly expressed in basal cells underlying the keratodonts and is seemingly absent from the lower jaw sheath (Fig. 5D), but papillae in the oral epithelium and overlying the infrarostral cartilage are positive for *Sox2*. These papillae likely house taste buds [33], and *Sox2* is a well-known taste-bud marker in vertebrates [33]. No *Shh* expression was detected in lower jaw oral epithelium, lower jaw keratinized sheath, or keratodonts.

**Fig. 5.**
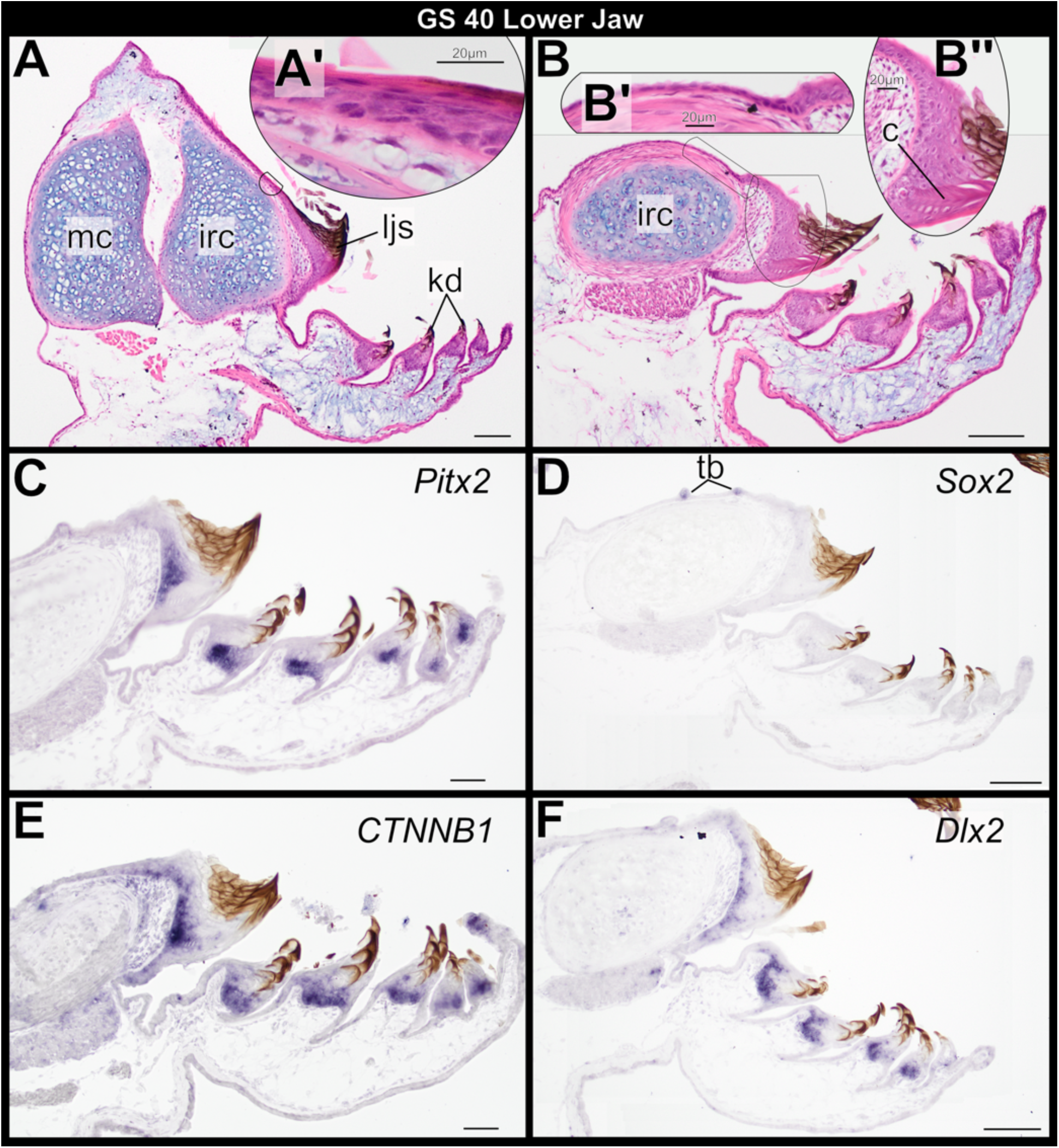
Sagittal sections through the larval lower jaw of *Osteopilus septentrionalis*, Gosner stage 40, stained with H&E (A, B) and *in situ* hybridization (C–F). Insets A’ and B’ show higher magnification detail of the lower jaw oral epithelium overlying the infrarostral cartilage; B” shows detail of the basal cells, sheath cells, and cone cells that form the jaw sheath. *Pitx2*, *CTNNB1*, and *Dlx2* are highly expressed in the tadpole jaw sheath and keratodonts (C, E, and F). *Sox2* is expressed in the larval taste buds overlying the infrarostral cartilage and in keratodont progenitor cells. Abbreviations: c, cone cells; irc, infrarostral cartilage; mc, Meckel’s cartilage; tb, taste buds. Scale bar, 100 µm.

Lower jaw histology of GS-41 specimens that retain keratinized mouthparts is very similar to that at GS 40, and there is no sign of tooth development (Fig. 6A). As the tadpole feeding apparatus atrophies during this stage, epithelium that previously supported the lower jaw sheath at the oral/aboral boundary retains columnar basal cells, which produce a thickened area (Fig. 6B and C). This thickening is broader than the localized epithelial thickenings that form in the upper jaw, it does not form a concave epithelial plate, and it does not persist into GS 42. We interpret it as a relic of the tadpole lower jaw sheath and unrelated to odontogenesis. Posteriorly, epithelium overlying the infrarostral cartilage transitions from single- to multi-layered as the tadpole feeding apparatus is lost (Fig. 6B and C), and oral epithelium of the lower jaw retains a stratified architecture for the remainder of metamorphosis (GS 42–46; Fig. 7A–H). One GS-42 specimen has an apparent epithelial thickening that invaginates slightly into the underlying mesenchyme (Fig. 7A). However, the basal cells of this thickening are not columnar, and the tissue architecture is inconsistent with formation of a dental lamina as seen in the upper jaw. We regard it as the remnant of a larval papilla or taste bud that is being resorbed, as other GS 41 and 42 specimens retain papillae in this general region that are in various stages of degeneration (Figs. 6B, 7C and D). There is no histological evidence of mandibular tooth formation in any GS 42–46 specimen.

**Fig. 6.**
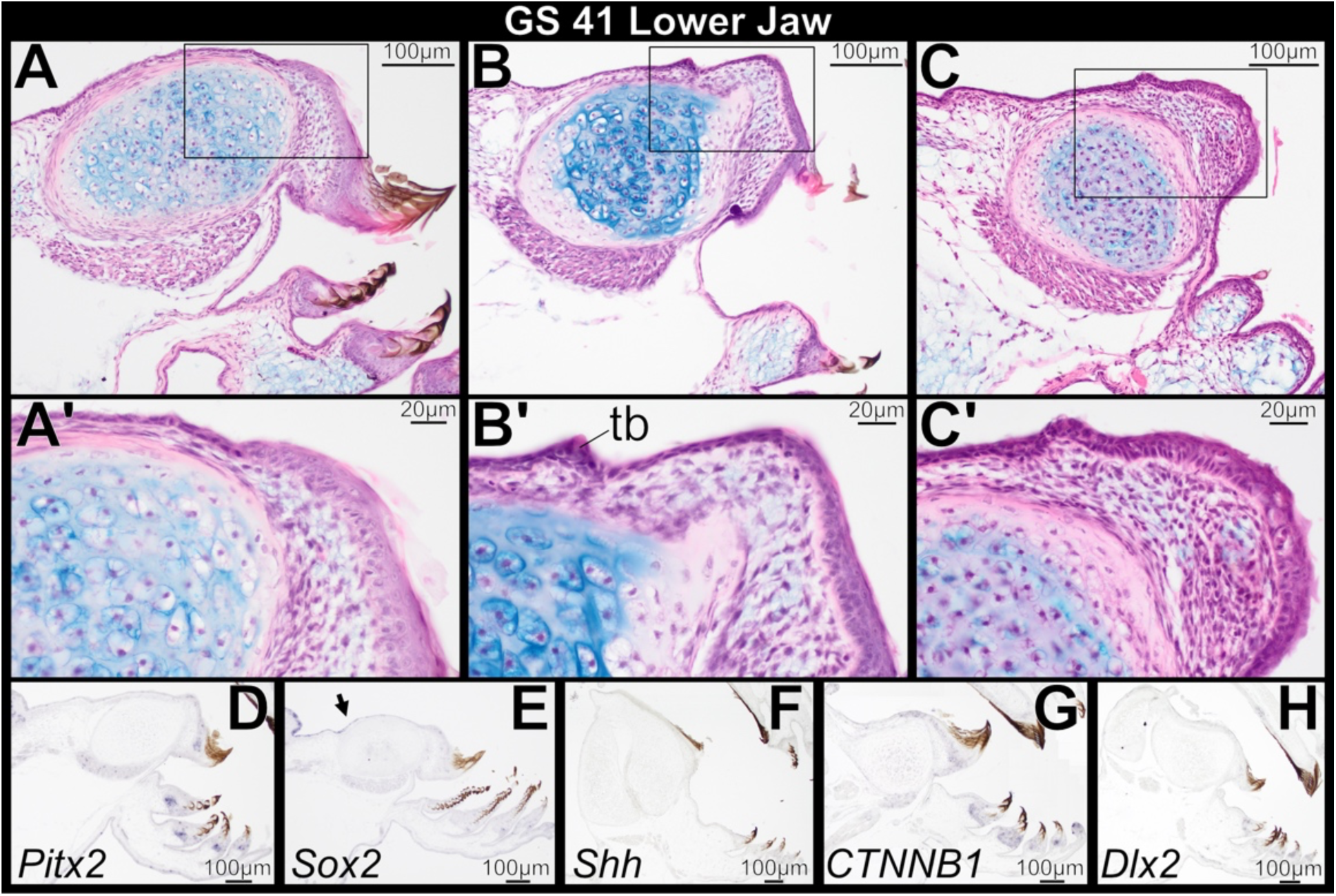
Sagittal sections through the larval lower jaw of *Osteopilus septentrionalis*, Gosner stage 41, stained with H&E (A–C) and *in situ* hybridization (D–H). Keratinized mouthparts degenerate (A, B), and columnar basal epithelial cells are a relic of the lower jaw sheath (C). Epithelium overlying the infrarostral cartilage transitions from single- to multi-layered. There is no histological evidence of mandibular tooth formation. *Pitx2*, *CTNNB1*, and *Dlx2* are weakly expressed in basal cells of the keratinized jaw sheath and keratodonts (D, G, and H). *Sox2* expression expands anteriorly from the tongue epithelium towards the oral epithelium (E, arrow).

**Fig. 7.**
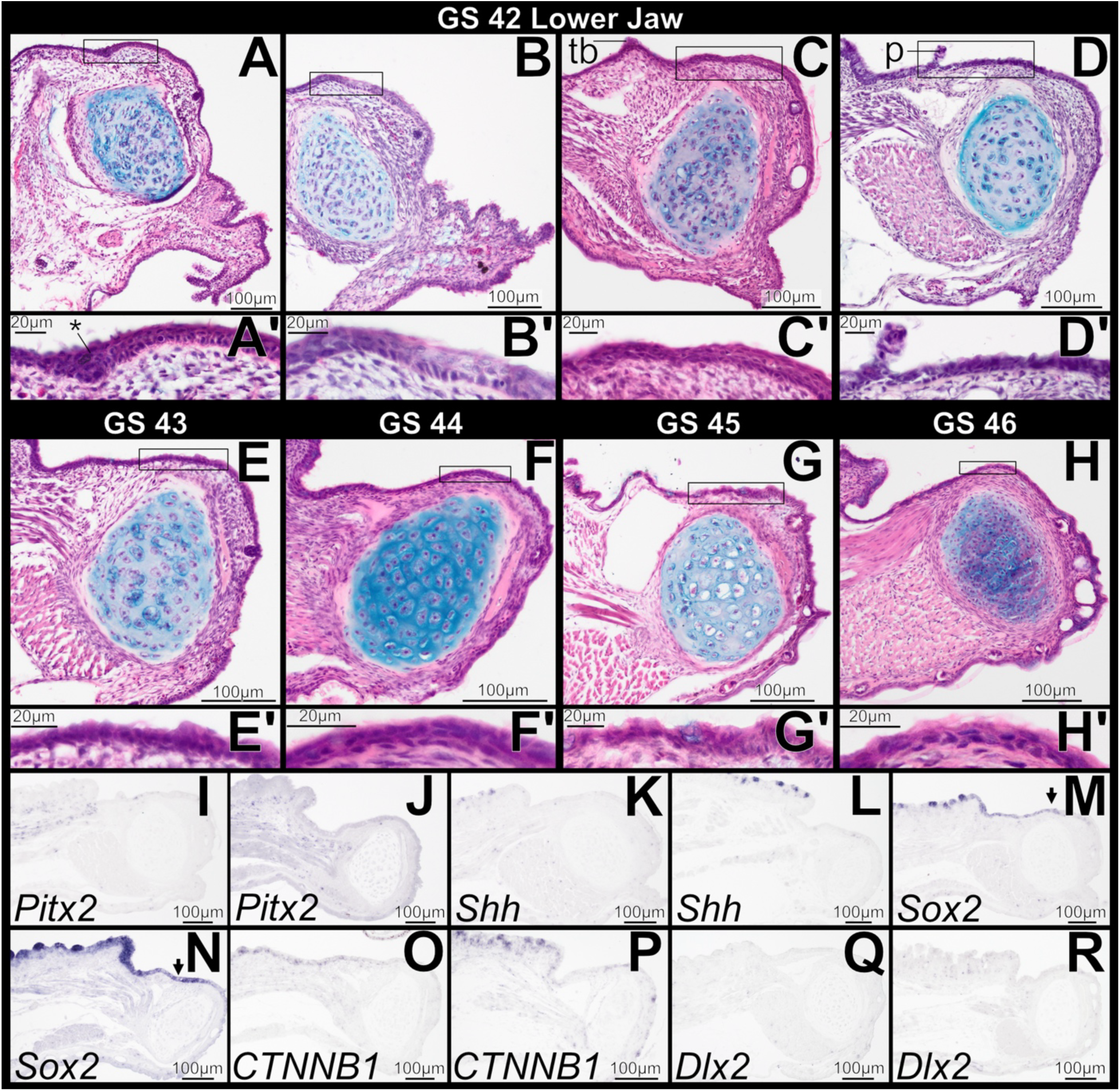
Sagittal sections through the lower jaw in *Osteopilus septentrionalis* during and immediately following metamorphosis, Gosner stages 42–46, stained with H&E (A–H) and *in situ* hybridization (I–R). Epithelium overlying lower jaw cartilage is multi-layered; there is no histological evidence of mandibular tooth formation. *Pitx2*, *Shh*, *CTNNB1*, and *Dlx2* is absent from mandibular oral epithelium (excluding tongue taste discs; I–L, O–R). *Sox2* is expressed in oral epithelium of the lower jaw (M and N, arrows). Abbreviations: p, papilla; *, invaginated epithelial thickening.

At GS 41, the genes *Pitx2*, *CTNNB1*, and *Dlx2* are all weakly expressed in basal cells of the keratinized jaw sheath and keratodonts prior to their atrophy (Fig. 6D, G, and H). These markers, however, are not expressed in the oral epithelium posterior to the jaw sheath and overlying the infrarostral, and they are not seen in the lower jaw at GS 42, following degeneration of tadpole mouthparts (Fig. 7I, O, and Q). Their absence persists through metamorphosis (Fig. 7J, P, and R). *Shh* is not expressed in the lower jaw between GS 41 and 46 (Figs. 6F, 7K and L). *Sox2* expression expands anteriorly from the lingual epithelium towards the epithelium overlying the infrarostral cartilage beginning at GS 41 (Fig. 6E), and by GS 42 it is seen within the oral epithelium of the lower jaw (Fig. 7M). It persists within the oral epithelium to later stages (Fig. 7N). *Shh*, *Sox2*, and *CTNNB1* are expressed in the developing taste buds of the tongue (Fig. 7L, N, and P).

### Formation of the primary mouth

We investigated gene expression patterns associated with stomodeum formation in embryos at GS 20 and 21 (Fig. 1A). These stages are characterized by rupture of the oropharyngeal membrane, which separates oral ectoderm from foregut endoderm [35–36], and formation of the primordial mouth. *Pitx2*, *Shh*, *Sox2*, *CTNNB1*, and *Dlx2* are all expressed within oral epithelium of the primordial upper and lower jaws (Fig. 8A–E). *Shh* and *Sox2* are limited to oral epithelium and foregut endoderm, whereas *Pitx2* and *Dlx2* have separate zones of expression in oral and aboral epithelium. *CTNNB1* is widely expressed in the epithelium and underlying mesenchyme of the primordial jaws.

**Fig. 8.**
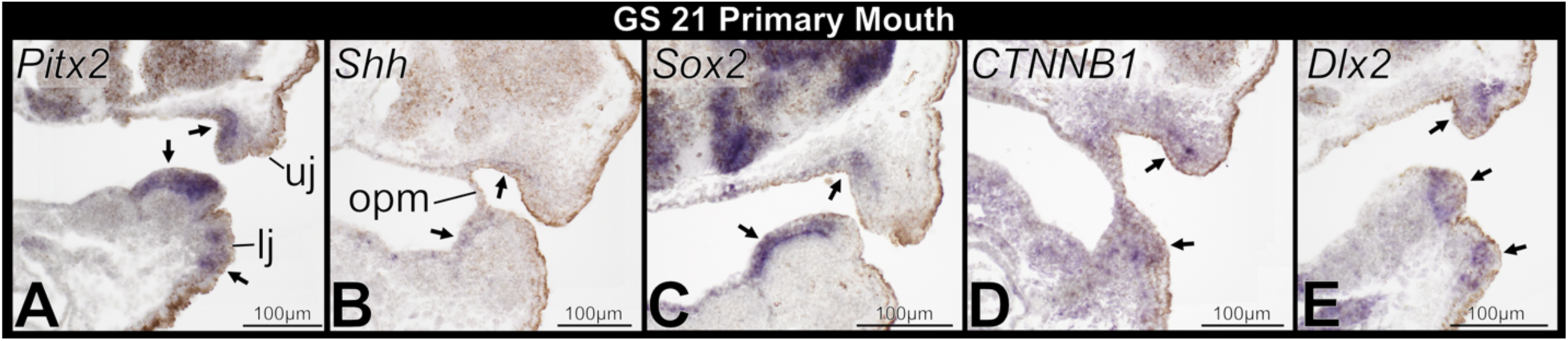
Sagittal sections through the primary mouth of embryonic *Osteopilus septentrionalis*, Gosner stage 21, following *in situ* hybridization. Arrows denote positive gene expression in the oral and aboral epithelium of upper and lower jaw primordia. Abbreviations: lj, lower jaw primordium; opm, oropharyngeal membrane; uj, upper jaw primordium.

## DISCUSSION

### Establishing dental competence and emergence of first-generation teeth of the frog upper jaw

Studies of turtles, birds, and non-avian theropod dinosaurs suggest that the presence of keratinized tissue overlying the jaw epithelia of vertebrates may in some instances dictate the location of tooth formation, and in others entirely prevent it [10, 37–38]. Our results with the Cuban tree frog support this hypothesis. Primary teeth begin to emerge on the upper jaw in *Osteopilus septentrionalis* before the keratinized tadpole mouthparts degenerate. However, presence of the upper jaw sheath appears to constrain the initial location of emerging tooth buds within the oral epithelium: the earliest epithelial thickenings and tooth placodes are positioned far from the oral/aboral boundary and posterior to the superficial, keratinized sheath cells that roof the anterior-most region of the oral mucosa (Figs. 2B, 3A–D). Following atrophy of the tadpole mouthparts and resorption of the suprarostral cartilage, additional tooth buds form closer to the tip of the upper jaw (Fig. 3).

The initiation of tooth development occurs in *X. laevis* and *X. tropicalis* at Nieuwkoop and Faber (NF) stages 55 and 56, respectively [8, 30]. These stages are equivalent to GS 35– 38 in *Osteopilus* and many other metamorphosing anurans [39] and precede metamorphic climax by several weeks. Some authors suggest that early onset of tooth formation in *Xenopus* compared to other frogs is due to the lack of keratinized mouthparts in this lineage [40], but we regard this as unlikely because *O. septentrionalis* possesses tooth buds and jaw sheaths concurrently. In the earliest description of anuran teeth, Hertwig [41] reported that tooth development in *Pelobates fuscus* begins at GS 42, following the loss of keratinized mouthparts. However, the tooth buds he describes are already at the cap stage of morphogenesis, well after onset of tooth development. The only other published histological studies of anuran odontogenesis examine replacement teeth that differentiate from the deeply invaginating successional dental lamina in adults [42–43]. More research is needed to establish the degree of variability in the timing of tooth induction across anuran diversity and determine if and how the formation of first-generation teeth is affected by the presence of keratinized tadpole mouthparts.

We tested for the expression of several key genes to assess if an odontogenic band (OB) precedes formation of teeth on the upper jaw in *O. septentrionalis*. The genes *Pitx2*, *Sox2*, and *Shh* are expressed in oral epithelium of the upper jaw at GS 40 (Fig. 2C–E), in the region where tooth placodes will form. Thus, frogs have dental precursor expression patterns consistent with an OB and comparable to those in chondrichthyans [3], teleosts [1, 34], salamanders [6], and amniotes [4, 44]. Our findings differ from those reported for *X. tropicalis*, in which *Shh* was not detected in upper jaw oral epithelium preceding tooth formation and was observed only in the inner dental epithelium of cap-stage tooth buds [8]. That study assessed gene expression by using wholemount *in situ* hybridization, whereas we examined serial sections. We show that *Shh* is weakly expressed in oral epithelium prior to tooth formation and during dental lamina emergence, but it is not more strongly expressed until the inner dental epithelium differentiates. Typically, *Pitx2*, *Shh*, and *Sox2* are all strongly expressed in the oral epithelium of vertebrates and are easily detected as a horseshoe-shaped band of expression in the jaws of whole-mount embryos [6, 44–46]. Frogs may be unique in having a less pronounced OB. Alternatively, wholemount *in situ* hybridization may be less effective in post-embryonic specimens, which have less-transparent tissues. The gene expression patterns we report in *O. septentrionalis* reaffirm that the core gene regulatory network underlying the initiation and morphogenesis of teeth is deeply conserved among vertebrates [3], including the late-forming teeth of anurans.

### Probing for the mechanism underlying the re-evolution of lost mandibular teeth

We find no definitive histological or genetic evidence of tooth development in the lower jaw of *O. septentrionalis*. The odontogenic competence of the lower jaw has mostly eroded. We looked for expression of several key genes to assess if a transient OB forms on the mandible, but we did not detect a zone of *Pitx2*, *Shh*, *CTNNB1*, or *Dlx2* expression in the oral epithelium overlying the lower jaw cartilages at any stage. *Sox2*, which regulates progenitor dental epithelial cells [4], is the only OB marker expressed within the mandibular oral epithelium, possibly indicating that the tooth GRN may not be completely lost. In cichlids, however, *Sox2* dramatically expands in the lower jaw following chemical knockdown of Bmp and Shh signaling, which ultimately yields fewer teeth [34]. Considering these results, it’s also possible that *Sox2* expression is expanded in the lower jaw of frogs due to the absence of other molecular signals within the oral epithelium.

The lack of tooth bud rudiments and absence of early odontogenic signaling in the lower jaw is unexpected because structures evolutionarily lost from adults typically form at least partly during development [24]. A transient odontogenic band, and sometimes vestigial tooth formation, is well documented in vertebrates that have lost teeth, including those in which teeth are entirely absent (e.g., edentulous mammals [47], birds [48–49], turtles [50–51]) and others in which tooth loss is spatially restricted (e.g., diastema teeth in rodents [44, 52]; oral jaw teeth in cypriniform fishes [53–54]). Frogs may be unique in failing to initiate the tooth development program within the lower jaw because of the dramatic decoupling between the formation of the primary mouth during embryogenesis and the onset of tooth development during metamorphosis. The presence of *Pitx2*, *Shh*, *Sox2*, *CTNNB1*, and *Dlx2* expression in the primordial oral epithelium of both upper and lower jaws during primary mouth formation in embryos of *O. septentrionalis* may indicate that these genes 1) are additionally involved in primary mouth formation, 2) are involved in the development of keratinized mouthparts, and/or 3) are involved in formation of an OB and establishment of epithelial competence to form teeth at an early embryonic stage, comparable to the timing of OB formation in other vertebrates. Shh and β-Catenin (*CTNNB1*) signaling are involved in primary mouth formation in embryonic *Xenopus* [55–56], but the role of these other markers in stomodeum development, if any, requires further investigation [36]. The genetics underlying keratinized mouthpart formation in frog embryos have not been studied previously, but these genes are likely involved in inducing jaw sheath and keratodont development based on their expression in late-stage tadpoles. One possibility is that some of the signaling pathways that typically orchestrate odontogenesis have been recruited to form keratinized mouthparts following primary mouth formation and are unable to participate in tooth development until jaw sheaths and keratodonts are no longer maintained. Further experimental work is needed in a comparative framework to evaluate the developmental genetics of the anuran mouth during primary mouth formation, development of the tadpole feeding apparatus, and transition to post-metamorphic jaws in diverse species to determine the interconnectivity and dynamics of these processes.

If the genes expressed during primary mouth formation in *O. septentrionalis* do not mediate formation of an OB and the early establishment of odontogenic potential, then the tooth development program may be entirely absent from the lower jaw of frogs. If this is the case, the re-evolution of mandibular teeth in *G. guentheri* will endure as one of the most perplexing exceptions to Dollo’s law of irreversibility. Further analysis of the mechanism underlying this extraordinary evolutionary reversal is constrained by the lack of living specimens and fresh tissues of this species, which is feared extinct [57]. Because *O. septentrionalis* and *G. guentheri* are somewhat distant relatives within the frog tree of life (same superfamily but different families), a promising avenue of research would be to investigate whether close relatives of *G. guentheri,* especially congeneric species, retain any odontogenic signaling in the lower jaw.

### Gene expression patterns overlap between tadpole keratinized mouthparts and frog teeth

Most vertebrates with a complex life cycle possess true teeth in both larval and adult forms (e.g., biphasic teleost fishes [58], extinct temnospondyl amphibians [59], salamanders and caecilians [9]). Frogs are therefore unique in that tadpoles possess a highly specialized feeding apparatus that lacks true teeth but is instead composed of keratinized mouthparts. The origin of the novel tadpole feeding apparatus is unknown, but the likely ancestral larval state in crown-group frogs is a tadpole with an oral disk that supports keratinized mouthparts [18]. If true, two key events occurred during the evolution of proto-anuran larvae: 1) tooth development was delayed from embryogenesis to metamorphosis, and 2) keratinized mouthparts originated and assumed the larval feeding function of teeth. The earliest morphological indication of keratinized mouthpart development occurs during embryogenesis, coincident with primary mouth formation [60]: thickening of the oral and aboral epithelium and condensation of underlying mesenchyme [61]. Unlike the invaginating epithelium of a tooth bud, developing jaw sheaths and keratodonts form as protruding outgrowths [61].

Keratinized mouthparts of tadpoles and true teeth of frogs are distinct with respect to cellular anatomy, tissue composition, and gross morphology [12]; shared genetic components should be unlikely. However, we observed an overlap in the complement of genes that are expressed in the jaw sheaths, keratodonts, and teeth of *O. septentrionalis*, indicating these structures may be more deeply related than previously recognized. Expression of *Pitx2* is particularly intriguing because this transcription factor is one of the most crucial regulators of the tooth GRN—the earliest epithelial marker of odontogenic competence [29, 62–63]. Furthermore, *Pitx2* is not regarded as a key developmental regulator of other keratinized ectodermal appendages (scales, hair, claws, etc.), although its overexpression in zebrafish embryos can lead to the formation of cutaneous “horn-like” structures, epidermal thickening, and increased expression of keratin genes [64]. Human epidermal keratinocytes can be induced to express *Pitx2* and differentiate into enamel-secreting ameloblasts when recombined with mouse embryonic dental mesenchyme [65]. Thus, *Pitx2*-expressing keratinocytes can form keratinized structures or alternatively gain odontogenic competence.

Our expression data are the first ever reported for keratinized mouthparts of tadpoles. Overlapping expression patterns between keratinized mouthparts and teeth may indicate that the core GRN underlying tooth formation was partially co-opted into the development of tadpole keratinized mouthparts during the early evolution of frogs. Co-option occurs during evolution when an ancestral gene or GRN that regulates the development of one structure becomes activated in a new developmental context [66–68] and gives rise to a different, non-homologous structure; it can play a key role in major morphological transitions [69]. For example, partial co-option of the limb/appendage signaling network is responsible for the origin and diversification of beetle horns [70] and butterfly wing eye spots [71]. Chimera experiments conducted nearly a century ago generated salamander larvae with keratinized mouthparts by transplanting frog-embryo ectoderm to the future mouth region of salamander embryos [72–74]. That such distinct organs can be induced by the same underlying ectomesenchyme signals [75] indicates some shared developmental program. Similar expression patterns between a novel structure and a preexisting organ can be a sign of GRN co-option, but functional experiments are needed to demonstrate that some of the common genes are functionally required for both traits [68]. Knockdown experiments of essential tooth GRN genes during tadpole mouthpart development may be a fruitful place to start.

Keratinized feeding structures have repeatedly evolved among vertebrates, including lampreys and hagfish [76], teleosts [77], frogs and sirenid salamanders [78], and amniotes [47]. In all cases except agnathans, keratinized structures evolved in the oral cavity to functionally replace true teeth that were evolutionarily lost. Expression patterns shared between teeth and tadpole keratinized mouthparts lend additional support to the hypothesis that the origin of keratinized feeding structures in vertebrates is developmentally linked to the tooth GRN in some way [47]. In birds, Bmp, Shh, Fgf, and Wnt signaling is involved in both odontogenesis and formation of the beak and rhamphotheca [79–80], and some of these signaling pathways may have been altered, inducing a diversion from tooth development to rhamphotheca formation [37]. Similarly, FGF4 protein signals are present during development of both tooth buds and baleen plates in the bowhead whale, and the tooth formation signaling cascade may have been co-opted to form baleen during the evolution of mysticete whales [81]. Additional characterization of keratin-to-tooth and tooth-to-keratin transitions among living vertebrates will shed light on the genomic and developmental features that may link these highly distinct yet seemingly related tissues.

## Conclusion

The postmetamorphic development of anurans continues to be enigmatic, and the unique contrast of dental conservation in the upper jaw coupled with complete loss of teeth in the mandible suggest deep rooted modules segregate the jaws. Further investigation is necessary to define this frog-specific evolutionary and developmental anomaly

## MATERIALS AND METHODS

### Specimen Sampling and Histology

We investigated tooth development in the Cuban tree frog, *Osteopilus septentrionalis*. Adults have a typical anuran dentition characterized by bicuspid, pedicellate teeth on the upper jaw and palate but an edentate lower jaw (Fig. 1; [82]). Tadpoles have a typical oral disk morphology, which includes keratinized upper and lower jaw sheaths and keratodonts (i.e., keratinized labial “teeth”) above and below the jaws. All tadpoles were collected from an invasive population in Gainesville, Alachua County, Florida (Florida Fish and Wildlife Conservation Commission permit number LSSC-12-00016C) and reared in captivity at the University of Florida (IACUC protocol #202111468). A developmental series across metamorphosis was obtained, ranging from Gosner developmental stages 40 to 46 (GS; [83]). In total, we investigated the histology of 45 specimens: seven GS 40, nine GS 41, nine GS 42, five GS 43, six GS 44, five GS 45, and four GS 46. Specimens were euthanized using buffered MS-222 and fixed in 4% paraformaldehyde at 4°C for 24–48 hr. Following fixation, specimens were dehydrated through a DEPC PBC-to-ethanol gradient and then stored in 100% ethanol at - 20°C until paraffin sectioning. Specimens were decalcified in 10% ethylenediaminetetraacetic acid (EDTA) for 48 hr, embedded in paraffin following standard protocols, and sectioned to 8–10 μm in sagittal or coronal planes using a rotary microtome at the University of Florida (Leica RM2145) or at the University of Dayton (Leica RM2155 or Rankin MCT25). Tissue sections were then either stained using a standard hematoxylin and eosin (H&E) plus Alcian blue protocol (SI Methods) to evaluate histological anatomy or processed for *in situ* hybridization to characterize gene expression patterns. Following examination of GS 40–46 specimens, we additionally obtained and sectioned multiple embryos to investigate gene expression patterns associated with primary mouth (i.e., stomodeum) formation during GS 20–21.

### RNA Probe Synthesis and Section in Situ Hybridization

We designed and synthesized RNA probes for five developmental genes with vital roles in the induction, early differentiation, and morphogenesis of teeth in fishes and amniotes [1–3]: *Sox2*, *Pitx2*, *Shh*, *CTNNB1*, and *Dlx2*. The OB is marked by expression of *Shh*, *Pitx2*, and *Sox2* [1–4], and both *CTNNB1* and *Dlx2* are often used as tooth bud markers [3, 53]. Digoxigenin-labeled antisense riboprobes were designed using sequence data available from frog genomes published on NCBI. Riboprobes were cloned using *O. septentrionalis* cDNA using the following primer sequences: ***Sox2*** (Forward GAGACCCATGAACGCCTTCATG; Reverse TAGTGCTGGGACATGTGCAG), ***Pitx2*** (Forward AGAGACAGAGGAGGCAGAGG; Reverse TGCCAGGCTGGAGTTACATG), ***Shh* (**Forward GACACCCTAAAAAGCTGACC; Reverse CAAATCCAGCCTCCACCGCCAG), ***CTNNB1*** (Forward AATTGCTGGTTCGTGCACAC; Reverse TCCCATTGCATCTTGCCCAT), and ***Dlx2*** (Forward ACCGGAGTGTTTGACAGCTTGG; Reverse TGTGATGATGGAGGTGGTGC). PCR clones were ligated into pGEM-T-Easy Vectors (Promega) and used as a template for probe synthesis. Digoxigenin-labeled antisense RNA probes were synthesized through in vitro transcription of the PCR templates with T7/SP6 RNA polymerase (Promega) and DIG RNA Labeling Mix (Roche). *In situ* hybridization was performed on the paraffin-embedded tissue sections using a standard protocol (SI Methods). *In situ* hybridization was performed multiple times for each probe to ensure that expression patterns were reproducible.

## Acknowledgements

This work was supported by the U.S. National Science Foundation (DBI-2109344 to DJP, DBI-2122620 to JH) and by the University of Dayton (UD) Department of Biology, a UD Research Council Seed Grant to DJP, and a UD College of Arts and Sciences Dean’s Summer Fellowship to MB, KG-B, and JS.

## Notes

### Competing Interest Statement

The authors have declared no competing interest.

