## Supplementary material for "Tooth development in frogs: Implications for the re-evolution of lost mandibular teeth and the origin of a morphological innovation": SI Methods

### Supporting Information Methods

#### Hematoxylin and Eosin (H&E) plus Alcian Blue Staining

Hematoxylin and eosin (H&E) plus Alcian blue staining was performed on the paraffin-embedded tissue sections using a standard protocol [1]. Tissue sections mounted on glass slides were deparaffinized with HistoClear II, rehydrated, and stained in 0.01% Alcian blue in 70% ethanol for 1 hr. Slides were then rinsed with tap water, stained in hematoxylin for 3 min, rinsed with tap water, dipped three times in 96% EtOH with 35% HCl, rinsed with tap water, and stained with eosin for 1.5 min. Following staining, the slides were washed with 96% ethanol for 8 min, 100% ethanol for 8 min, and HistoClear II for 8 min. Sections were mounted using DPX mountant, covered with a glass coverslip, and imaged using an Olympus BX53 LED compound light microscope and Olympus DP23 camera.

#### Section in Situ Hybridization

*In situ* hybridization was performed on the paraffin-embedded tissue sections using a standard protocol [2]. Tissue sections mounted on glass slides were deparaffinized with HistoClear II, rehydrated, and then incubated in prehybridization buffer for 2 hr. All DIG-labeled antisense RNA probes were denatured at 95°C for 12 min immediately before being applied to the tissue sections, which were then incubated overnight at 61°C. The following day, the slides were washed in 2× SSC and 0.2× SSC at 61°C and then incubated in blocking solution (2% Roche Blocking Reagent in maleic acid buffer containing Tween 20 [MABT]) for 1 hr at room temperature. Slides were antibody-labeled overnight at 4°C with anti-DIG-ALP (0.5 µL/mL; Roche). Following six MABT washes, substrate color reactions were carried out by adding 300 µL of BM purple (Roche) onto each slide at room temperature and replenished every 24 hr until the reactions were sufficient to visualize gene expression but minimized background signal. Slides were then fixed for 1 min in 4% PFA, washed in PBS and distilled water, mounted with Fluoromount-G with DAPI (Thermo Fisher), and imaged using an Olympus BX53 LED compound light microscope and Olympus DP23 camera. *In situ* hybridization was performed multiple times for each probe to ensure that expression patterns were reproducible.

#### Computed Tomography Scanning

We performed high-resolution X-ray computed tomography (CT) scanning using a GE v|tome|x M 240 system at the Nanoscale Research Facility (University of Florida, Gainesville, USA). We conducted scans using a 180 kv X-ray tube containing a diamond-tungsten target, with the voltage and the current adjusted to maximize absorption range for each specimen. We processed the raw X-ray data using GE's datos|x v2.3 software, producing tomogram and volume files. We imported these microCT volume files into VG StudioMax v3.5 (Volume Graphics, Heidelberg, Germany) and isolated regions of interest using the suite of segmentation tools in VG StudioMax. We used diffusible iodine-based contrast-enhanced CT [3] to image Gosner stage 40 and 42 tadpoles of *Osteopilus septentrionalis* from lot UF-H-184921. Tadpoles were soaked in 1.25% $I_2KI$ (Lugol's solution) for 72 hours prior to scanning. We CT scanned an unstained adult *Osteopilus septentrionalis* specimen (UF-H-63656). We deposited image stacks (TIFF) in MorphoSource (tadpoles: https://www.morphosource.org/concern/biological_specimens/000706050/ [DOIs requested]; adult: https://doi.org/10.17602/M2/M34831).
