## Supplementary material for "Tooth development in frogs: Implications for the re-evolution of lost mandibular teeth and the origin of a morphological innovation": Fig. S1

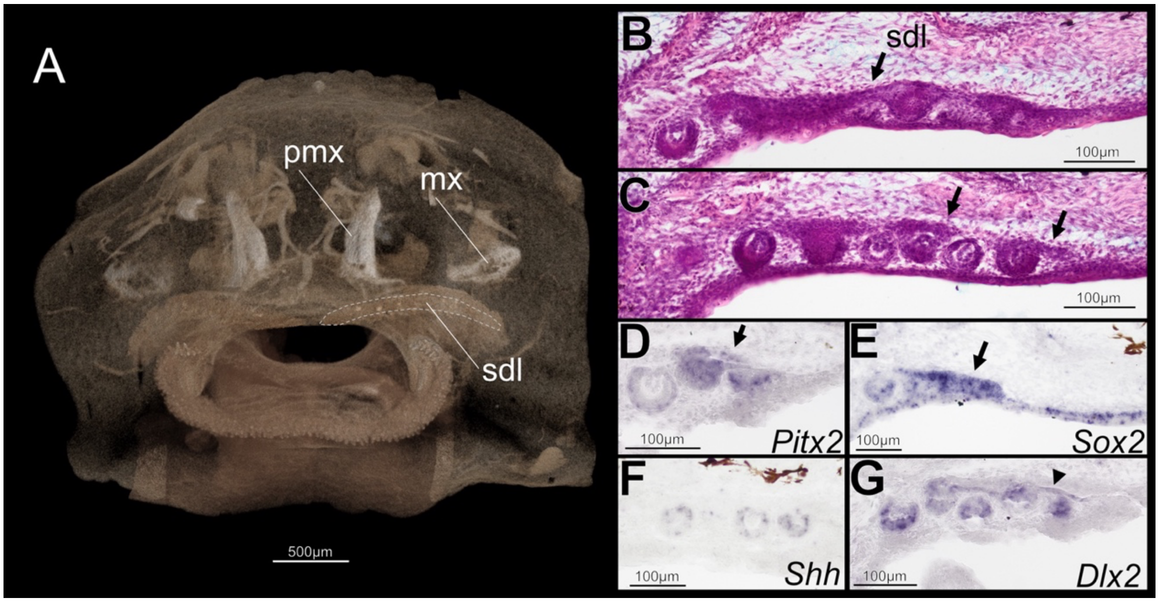


**Fig. S1.** Successional dental lamina in the upper jaw of *Osteopilus septentrionalis*, Gosner stage 42, visualized by microCT (A), coronal sections stained with H&E (B, C) and *in situ* hybridization (D–G). The successional dental lamina (arrows) adjoins the row of emerging tooth buds and is marked by *Pitx2*, *Sox2*, and *Dlx2* expression (D, E, and G). *Shh* is expressed in the inner dental epithelium of tooth buds but not in the dental lamina (F). Abbreviations: mx, maxilla; pmx, premaxilla; sdl, successional dental lamina.
